# An Fc-silent OspA monoclonal antibody passively protects mice from tick and intradermal *Borrelia burgdorferi* challenge

**DOI:** 10.64898/2025.12.14.694235

**Authors:** Daniel Palmer, Atieh Shemshadian, Katherine Berman, Graham G. Willsey, Carol Lyn Piazza, Grace Freeman-Gallant, Michael J Rudolph, Jeff Bourgeois, Linden Hu, David J. Vance, Nicholas Mantis

## Abstract

The monoclonal antibody, LA-2, has played a pivotal role in the development of <u>O</u>uter <u>s</u>urface <u>p</u>rotein <u>A</u> (OspA)-based vaccines for Lyme disease, a multisystem illness caused by the tick-borne spirochete, *Borrelia burgdorferi* sensu lato. Of particular significance was the demonstration more than three decades ago that LA-2 equivalent antibody titers, defined by a competitive-inhibition ELISA, serve as a reliable correlate of vaccine-induced protection across different species, including humans. *In vitro* characterization of LA-2 has identified both complement-dependent and -independent activities, although which of these attributes contribute to protection against *B. burgdorferi* remains unresolved. To address this issue, we generated and characterized an “Fc-silent” version of LA-2 IgG1 carrying so-called LALAPG substitutions (L234A, L235A, P329G). We demonstrate that LA-2 LALAPG retained OspA binding activity but was severely attenuated in *in vitro* complement deposition and complement-dependent borreliacidal assays. Nonetheless, LA-2 LALAPG was as effective as LA-2 at passively protecting C3H mice against nymphal tick-mediated *B. burgdorferi* challenge. LA-2 LALAPG was also equivalent to LA-2 in passively protecting BALB/c mice against intradermal *B. burgdorferi* challenge. In the intradermal challenge model, viable spirochetes were not recoverable 24 h after injection from skin biopsies of mice treated with LA-2 or LA-2 LALAPG, and an influx of pro-inflammatory cytokines and chemokines to the injection site was abrogated. Collectively, these results suggest that LA-2’s primary mode of action involves direct physical interactions with the spirochete rather than complement-dependent killing. Elucidating these mechanisms may have implications for understanding the mechanistic correlates of OspA-based vaccine-induced immunity in humans.

## INTRODUCTION

Over the past five decades, monoclonal antibodies (mAbs) have emerged as extraordinary tools in the identification of protective antigens and epitopes associated with pathogens of interest, leading to novel vaccines for viruses, bacteria and parasites ^1–3^. In the case of the Lyme disease spirochete, *Borrelia burgdorferi* sensu lato, multiple groups working in the 1990s generated large collections of mAbs that led to the identification Outer surface protein A (OspA) as a candidate Lyme disease vaccine antigen ^4–15^. OspA is a lipoprotein expressed at high levels by *B. burgdorferi* within the midgut of its arthropod vector, the black legged tick (*Ixodes scapularis*), where it is proposed to function as an adhesin ^16^. Structurally, OspA consists of 21 anti-parallel β-strands with a single C-terminal α-helix ^17,18^. The N-terminus is anchored in the spirochete outer membrane via a lipid moiety, while the C-terminus projects away (∼80 Å) from the bacterial surface and is accessible to antibody attack ^19,20^. OspA is downregulated during or just after spirochete transmission to a mammalian host ^21,22^. As such, antibodies elicited by OspA-based vaccines are proposed to inhibit one or more steps in *B. burgdorferi* tick-mediated transmission, although the specific mechanisms by which this occurs remains to be fully elucidated.

Among the many OspA mAbs characterized to date, LA-2 has played a particularly significant role in our understanding of OspA-mediated immunity. LA-2 was one of the first OspA-specific mAbs shown to passively protect mice from *B. burgdorferi* needle infection ^6^ and tick-mediated challenge ^23^. And, until just a few years ago, LA-2 was the only protective antibody whose epitope on OspA had been resolved at the structural level ^18,24,25^. In the context of Lyme disease vaccine development, LA-2 serological antibody “equivalence,” as defined by a competitive ELISA, proved to correlate with protection against tick-mediated *B. burgdorferi* infection in OspA-vaccinated mice and dogs ^26^. Remarkably, as part of a large randomized OspA vaccine trial, it was determined in a subset of individuals that LA-2 equivalent titers are also important biomarkers of Lyme disease susceptibility in humans, as individuals with confirmed Lyme disease had lower LA-2 equivalence than those who did not ^27,28^. LA-2 continues to be used as a benchmark in the development of next generation OspA vaccines ^29^ (M. Finn, personal communication).

Despite LA-2’s central role in Lyme disease vaccine development, the exact mechanism by which LA-2 protects against *B. burgdorferi* infection remains incompletely defined. In fact, the basic question of whether complement is needed for LA-2’s protective activity has not been addressed. While LA-2 has potent complement-dependent borreliacidal activity *in vitro* ^23,30,31^, evidence indicates that complement (human or mouse) is not active in the tick midgut ^32^. Several complement-independent activities have been ascribed to LA-2, including effects on spirochete transmigration, that would be expected to impede *B. burgdorferi* from the tick midgut ^33,34^. Defining the contribution of complement in LA-2’s mechanism of action is important for understanding correlates of OspA-mediated immunity, especially as clinical trials of next generation OspA vaccines are ongoing ^29,35,36^. In this report, we generate an “Fc-silent” version of LA-2 IgG1 that is effectively devoid of *in vitro* complement-dependent borreliacidal activity and characterize its activity in mouse models of tick-mediated and intradermal *B. burgdorferi* challenge.

## Materials and Methods

### Ethics statement

The mouse experiments described in this study were reviewed and approved by the Institutional Animal Care and Use Committees (IACUC) at the Wadsworth Center (protocol 23-459) and Tufts University-Tufts Medical Center (protocol B2024-50). The Wadsworth Center and Tufts University-Tufts Medical Center both comply with the Public Health Service Policy on Humane Care and Use of Laboratory Animals and were issued assurance numbers A3183-01 and A4059-01, respectively. Both facilities are fully accredited by the Association for Assessment and Accreditation of Laboratory Animal Care (AAALAC). Obtaining this voluntary accreditation status reflects that these facilities’ Animal Care and Use Program meets all standards required by law and goes beyond the standards as it strives to achieve excellence in animal care and use. All animals were euthanized by carbon dioxide asphyxiation followed by cervical dislocation, as recommended by the Office of Laboratory Animal Welfare (OLAW), National Institutes of Health.

### Recombinant *B. burgdorferi* B31 proteins

Recombinant OspA, DbpA and OspC type A derived from *B. burgdorferi* strain B31 (**Table 1**) were expressed in *E. coli* as cited in **Table 1**.

**Table 1.** Recombinant *B. burgdorferi* B31 proteins used in this study.

| Table 1. Recombinant *B. burgdorferi* B31 proteins used in this study |  |  |  |
| --- | --- | --- | --- |
| Antigen | AA | UniProt ID | References |
| OspA | 18-273 | P0CL66 | 24 |
| OspB | 50-296 | P17739 | unpublished |
| OspC<sub>A</sub> | 38-201 | Q07337 | 37 |
| DbpA | 26-188 | O50917 | 38 |
| DbpB | 21-187 | O50917 | unpublished |
| Abbreviations; AA, amino acid; UniProt (https://www.uniprot.org/) |  |  |  |

### LA-2 and LA-2 LALAPG IgG1 expression and purification

Codon optimized V_H_ and V_L_ DNA sequences of LA-2 (Antibody Registry RRID: AB_2619693) derived from PDB 1FJ1 ^18^ were custom synthesized by Life Technologies (San Diego, CA) and cloned into TMV and PVX plant expression vectors containing codon-optimized human kappa and human IgG1 constant regions ^39^. The resulting plasmids were transformed into *Agrobacterium tumefaciens*. Four-week-old *N. benthamiana* plants were infiltrated with *A. tumefaciens* carrying plasmids for the expression of heavy and light chains of LA-2 LALAPG. Aerial plant parts were harvested after 7 days and extracted and clarified. The LA-2 LALAPG antibody was then purified with Protein A affinity and anion exchange chromatography ^40^.

### Antibody affinity determinations by Biolayer interferometry (BLI)

Affinity determinations were conducted using an Octet RED96e Biolayer Interferometer (Sartorius, Goettingen, Germany) with Data Acquisition 12.0 software. Biotinylated OspA (5 μg/mL) in PBS containing 2% w/v BSA (“buffer”) was captured onto Octet SA (streptavidin) biosensors (Sartorius) for 5 min. After equilibration, sensors were immersed in two-fold serial dilutions of mAb starting at 100 nM for 10 min. The sensors were then dipped into buffer for 30 min to allow for dissociation. The raw sensor data were loaded into the Data Analysis HT 12.0 software, grouped and fit using a 1:2 bivalent analyte model.

### Flow cytometric analysis of *B. burgdorferi* surface labeling

*B. burgdorferi* strain B31 surface labeling with LA-2 and LA-2 LALAPG was performed essentially as described ^31^. LA-2 and LA-2 LALAPG were 2-fold serially diluted in PBS before incubation with viable *B. burgdorferi* B31. The ricin-specific mAb, PB10, was used as an IgG1 isotype control (10 µg/mL). Alexa Fluor 647-labeled goat anti-human IgG (H+L) (Invitrogen, Carlsbad, CA) was used as a secondary antibody. Samples were analyzed using a BD FACSCalibur (BD Biosciences, Franklin Lakes, NJ). Bacteria were gated on FSC and SSC to exclude debris, and 20,000 events were counted per condition. Agglutination was calculated as the percent of events in the UL + UR + LR quadrants. Data were analyzed using FlowJo v10.10.0 (BD Biosciences).

### Complement-dependent borreliacidal assays

Complement-dependent borreliacidal assays were performed using a recombinant *B. burgdorferi* B31-5A4 strain carrying an IPTG-inducible *mscarlet-I* reporter (GGW979), as described ^31^. Briefly, GGW979 cultures were grown to mid-log phase in BSKII medium supplemented with gentamicin (50 µg/ml) at 32 °C under static conditions. Spirochetes were harvested by low-speed centrifugation and resuspended in phenol red–free BSKII containing gentamicin (50 µg/ml) to a final density of 3×10⁷ spirochetes/ml.

Cell suspensions were then mixed 1:1 with phenol red–free BSKII supplemented with 20% guinea pig complement (Sigma Aldrich, St. Louis, MO) and 20 nM of one of the following mAbs: LA-2, LA-2 LALAPG, 857-2, PB10. PB10, a ricin toxin-specific antibody, was used as an IgG1 isotype control. Reactions were set up in white 96-well assay plates (Co-Star). Following sample addition, reactions contained 1.5×10⁶ spirochetes, 5 nM of antibody and 10% guinea pig complement. Assay plates were then incubated overnight in a water jacketed incubator at 37°C with 5% CO_2_. The following day, 1 mM IPTG was added to each well to induce *mScarlet-I* expression. Following a 48-h incubation at 37°C with 5% CO_2_, the MFI was recorded at 569 nm (excitation)/611 nm (emission) using a Spectramax ID3 plate reader (Molecular Biosystems, San Jose, CA). Raw MFI data was then normalized as described ^31^. The data presented is the mean and SD of three independent experiments.

### MIA

*B. burgdorferi* B31 antigens OspA, OspB, OspC, DbpA, and DbpB (**Table 1**) were coupled to Magplex-C microspheres (5 μg antigen/ 1×10^6^ microspheres) using a xMap Antibody Coupling Kit as recommended by the manufacturer (Luminex Corporation, Austin, TX). Beads were protected from light and stored at 2-8°C in xMAP AbC Wash Buffer (5×10^6^ microspheres/mL) until use. Serum samples (1:100) and coupled microsphere stocks (1:50) were diluted in assay buffer (1 x PBS, 2% BSA, pH 7.4). The diluted sera (50 μL) and diluted microspheres (50 μL) were combined in black, clear-bottomed, non-binding, chimney 96-well plates (Greiner Bio-One, Monroe, North Carolina) and incubated at room temperature for 1 hr in a tabletop shaker (600 rpm). Plates were placed on a magnetic separator and washed three times using wash buffer (1 x PBS, 2% BSA, 0.02% TWEEN-20, 0.05% Sodium azide, pH 7.4). To detect seroconversion in the mice, goat anti-mouse IgG, Human-ads-PE (SouthernBiotech, Birmingham, Alabama) secondary antibody was diluted 1:500 in assay buffer, added (100 μL) to each well, and allowed to incubate at room temperature for 30 min in a tabletop shaker (600 rpm). Alternatively, to detect remaining LA-2 or LA-2 LALAPG, PE labeled goat anti-Human IgG Fc, eBioscience (Invitrogen) secondary antibody was used. Plates were washed as previously stated. The microspheres were resuspended in 100 μL of wash buffer and placed back on the tabletop shaker (600 rpm) for 5 minutes prior to analysis using a FlexMap 3D (Luminex Corporation). To establish reactivity cutoffs for each antigen, the average median fluorescent intensity (MFI) of buffer-only wells was multiplied by six. MFIs for each mouse serum sample were divided by the antigen-specific reactivity cutoffs yielding an index value. An index value greater than 1 suggests reactivity above background for the given antigen.

### Antibody-dependent complement deposition (ADCD) assay

We modified a flow cytometry-based HIV-1 antibody-dependent complement deposition (ADCD) for use with a Luminex instrument and OspA-coupled beads ^41^. Magplex-C microspheres coupled with recombinant *B. burgdorferi* antigens, OspA and OspC_A_, were diluted (1:50) and mixed 1: 1 (v/v) with primary antibodies, LA-2 and LA-2 LALAPG (10 μg/mL), then seeded into a 96-well plates, covered in foil, and incubated for 1 h at room temperature (RT) with shaking. Plates were washed twice using a plate magnet and 190 μL of wash buffer (PBS, 2% BSA, 0.02% Tween-20, 0.05% sodium azide, pH 7.4). Following the washes, 200 μL of diluted human complement (1:50; Pel-Freez Biologicals, Rogers, AR) were added to each well and incubated for 20 min at RT with shaking. Plates were washed again then phycoerythrin-tagged mouse anti-C3/C3b/iC3b (1:100; BD) and phycoerythrin-tagged goat anti-human IgG Fc (1:500; Invitrogen) were added to their respected wells and incubated for 30 min. The plates were washed a final time before antibody-bead complexes were resuspended in 100 uL of wash buffer and incubated for 1 min while shaking. The plates were analyzed via a FlexMAP 3D instrument (Luminex Corporation) with results presented in median fluorescence intensity (MFI).

### Mouse model of *B. burgdorferi* challenge by *Ixodes scapularis* nymphs

Animal studies were conducted with approval by the Institutional Animal Care and Use Committees (IACUC) at the Wadsworth Center and Tufts University-Tufts Medical Center. To generate *B. burgdorferi* B31 infected *Ixodes scapularis* nymphs, C57BL/6J mice were injected subcutaneously with 10^5 cells/mL log growth phase of *B. burgdorferi* B31. Two to four weeks later, the mice were infested with naive *Ixodes scapularis* larvae, which were allowed to feed to repletion. Replete larvae were harvested and allowed molt into mature infected nymphs. As controls, we used naive *Ixodes scapularis* nymphs procured from the Oklahoma State University tick rearing facility ^42^.

For challenge studies, equal numbers of male and female C3H/HeN mice aged ∼6 weeks (Charles River Laboratories, Kingston, NY) were acclimated in the Wadsworth Center’s vivarium for 1-2 weeks before the start of the experiment. On study day -1, mice were subcutaneously (SC) administered either LA-2, LA-2 LALAPG or an IgG1 isotype control (anti-*Vibrio cholerae* mAb ZAC-3) (120 μg or 30 μg per mouse) diluted in 200 µL of PBS. The following day, infected or naive nymphal ticks (5 per mouse) were placed on a shaved area of the mouse’s dorsum. Nymphs were collected from all mice 3-5 days post placement. Mice which had at least one tick that appeared to be at or near repletion at the time of collection were presumed to be successfully challenged. On study day 21, mice were euthanized, and blood was collected via cardiac puncture for serological analysis. Bladders were also collected for cultivation of spirochetes in 2 mL BSKII cultures treated with rifampicin (50 μg/mL), fosfomycin (20 μg/mL), and amphotericin B (2.5 μg/mL). Infection status was based on seroconversion using the MIA described above, as well as the presence or absence of live spirochetes in bladder cultures using dark-field microscopy, which were assessed weekly for one month.

### Collection of engorged ticks, dissection, and determination of genome equivalents

One replete or near-replete nymph that had fed on each mouse was dissected to harvest the midgut tissues. Midgut tissues were extracted using the E.Z.N.A.® Mollusc & Insect DNA Kit (Omega Bio-tek, Inc., Norcross, GA) and real-time qPCR was performed to determine the *Borrelia* burden in midgut tissues. The single-copy *B. burgdorferi flaB* (flagellin) gene was amplified, and *flaB* copy number was standardized to total gDNA in each sample as measured using the Qubit 4 Fluorometer (Invitrogen) to determine normalized spirochete burdens in tick midguts.

### Mouse model of intradermal *B. burgdorferi* challenge

Female BALB/c mice aged ∼8 weeks (Taconic Biosciences, Germantown, NY) were acclimated in the Wadsworth Center’s vivarium for one week before the start of the experiment. On study day -1, mice were injected intraperitoneally (IP) with LA-2 or LA-2 LALAPG (0.1 - 120 μg/mouse) in 200 µL PBS. The following day (study day 0), mice were challenged with mid-log phase *B. burgdorferi* strain B31-5A4 (1×10^5^ cells) by intradermal (ID) injection. On day 21, the mice were euthanized, and blood was collected via cardiac puncture for serological analysis. Infection status was determined based on seroconversion using the MIA described above.

To assess the effects of LA-2 and LA-2 LALAPG on *B. burgdorferi* skin dissemination, a mix of male and female BALB/c mice aged ∼6 weeks were injected SC with 30 µg of LA-2 or LA-2 LALAPG in 200 µL PBS, or remained untreated. The following day (study day 0), mice were challenged with mid-log phase *B. burgdorferi* strain B31-5A4 (1×10^5^ cells) by ID injection. Groups of mice were euthanized on days 1, 3 and 7, and ∼5 mm skin biopsies were excised from the injection site (IS), ∼1 cm away from the IS, and ∼3 cm away from the IS. Biopsies were rinsed in PBS immediately after collection and placed in 2 mL BSKII medium supplemented with rifampicin (50 μg/mL), fosfomycin (20 μg/mL), and amphotericin B (2.5 μg/mL) for cultivation of spirochetes. The biopsy cultures were assessed weekly by dark-field microscopy over the course of four weeks for the presence of viable spirochetes.

### Inflammatory cytokine and chemokine analysis in mouse skin biopsies

Groups of male and female BALB/c mice were SC administered 30 ug LA-2 per mouse on day -1 or left untreated. The following day (study day 0), mice were challenged with mid-log phase B. burgdorferi strain B31-5A4 (1×10^5^ cells) by ID injection. 5 days-post infection, mice were euthanized, and a skin biopsy ∼1 cm in diameter was collected from the injection site of each mouse in a cytokine extraction buffer containing 0.4M NaCl, 0.05% Tween 20, 0.5% Bovine Serum Albumin, 0.1 mM phenylmethylsulphonyl fluoride, and 20 Ki of aprotinin in 1X PBS. The solution containing the biopsy was homogenized at 5 m/s in a bead beater in 1-minute increments, with a 1-minute cool down between shakes. This was repeated five times, or until the biopsy was fully homogenized. This solution was centrifuged at 13,000 g for 10 minutes at 4°C, and the supernatant was collected and then prepared for cytometric bead array analysis. Skin homogenates were diluted 1:2 in assay diluent and processed using the BD Biosciences Cytometric Bead Array (CBA) Mouse Inflammation Kit following the manufacturer’s instructions. Samples were analyzed using a BD FACSCalibur (BD Biosciences).

### Statistical analysis

Statistical procedures for experiments are described in the figure legends. All statistical analysis of data was performed in R, Graphpad Prism 9.0, and Microsoft Excel. In all experiments, p-values <0.05 are considered significant.

## Results

### Recognition of recombinant and native OspA by LA-2 and LA-2 LALAPG

To generate an Fc-silent version of LA-2, codon optimized DNA sequences encoding the variable heavy chain (V_H_) was cloned in-frame into human IgG1 Fc and IgG1 LALAPG expression vectors. The IgG1 LALAPG derivative carries three-point mutations (L234A, L235A, P329G) relative to IgG1 that abolishes complement fixation activity and FcγR recognition ^43,44^. The LA-2 V_L_ coding sequence was inserted into a human kappa expression vector. The V_H_ and V_L_ plasmids were transformed into *A. tumefaciens* that was then used to infiltrate *N. benthamiana*. Aerial plant parts were harvested after 7 days and extracted and clarified antibodies were purified to homogeneity by Protein A affinity and anion exchange chromatography ^40^. By flow cytometry, LA-2 IgG1 and LA-2 LALAPG were equivalent in their ability to recognize native OspA on the surface of viable *B. burgdorferi* strain B31, as well as induce agglutination of those cells (**Figure 1**). Surface labeling was dose-dependent and resulted in a maximum of ∼90% total cell labeling with a median fluorescence intensity (MFIs) exceeding 7500 for LA-2 LALAPG. LA-2 and LA-2 LALAPG also recognized recombinant OspA with similar apparent affinities as measured by BLI (**Figure S1**).

**Figure 1.**
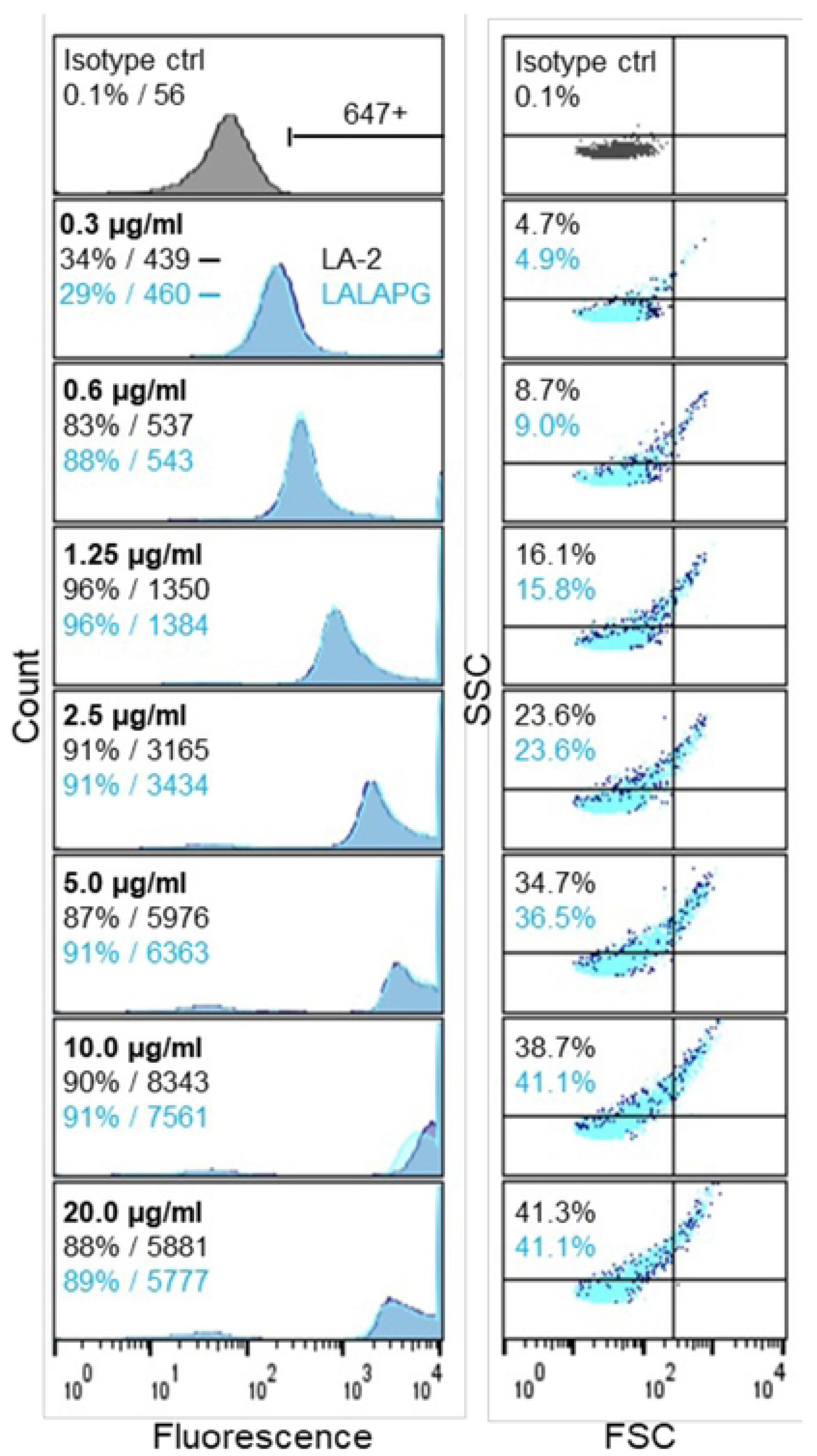
Reactivity of LA-2 and LA-2 LALAPG with native OspA. Representative flow cytometry assay of serially diluted LA-2 (dark blue) and LA-2 LALAPG (light blue overlay) reactivity with native OspA on the surface of *B. burgdorferi* B31. (Left) Fluorescence histogram overlays comparing binding properties. Percent and gMFIs of Alexa-647 fluorescently labeled events (under bracket) are indicated. (Right) Forward scatter (FSC) – side scatter (SSC) dot plot overlays comparing agglutination properties. Events increased in FSC and/or SSC (UL, UR, LR quadrants) demonstrate agglutination, and percent of events agglutinated is indicated.

### LA-2 LALAPG IgG1 lacks complement fixation and borreliacidal activities

Introduction of the LALAPG (L234A, L235A, P329G) mutations into the Fc region of IgG1 is reported to essentially eliminate complement fixation activity ^43,44^. To examine this in the case of LA-2, we adopted an antibody-dependent complement deposition (ADCD) assay to OspA (**Figure 2A**) ^41^. Recombinant OspA was covalently coupled to fluorescent microspheres, then probed with LA-2 IgG1 and LA-2 LALAPG in the presence of human complement. Total mAb binding to the beads was determined using PE-labeled anti-human IgG, while complement deposition was measured using a PE-labeled anti-C3 antibody. The results confirmed that LA-2 and LA-2 LALAPG have equivalent capacities to bind OspA (**Figure 2B**). However, the two mAbs were starkly different in terms of complement fixation activity. LA-2 demonstrated a dose-dependent increase in C3 deposition that peaked at ∼2 μg/ml. LA-2 LALAPG, in contrast, was devoid of any activity even at 10 μg/ml (**Figure 2C**). These results confirmed that LA-2 LALAPG is unable to fix complement via the classical pathway.

**Figure 2.**
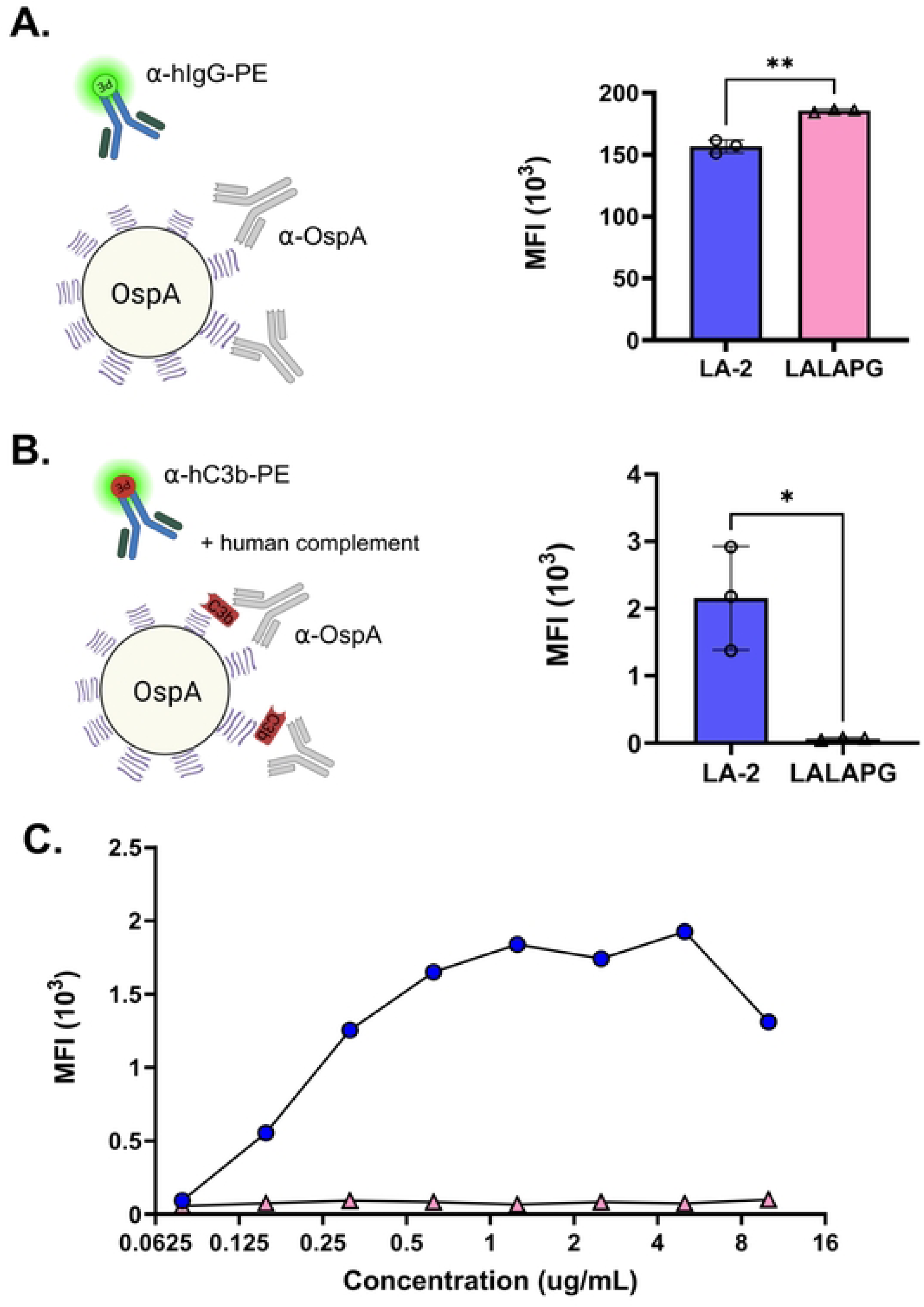
LA-2 LALAPG is deficient in complement deposition *in vitro*. (A) Binding (MFI) of LA-2 IgG and LA-2 LALAPG to recombinant OspA. The asterisks indicate a significant difference between groups by Welch’s t-test (**P<0.01). Quantification and comparison of human complement C3 deposition between LA-2 and LA-2 LALAPG in the context of OspA. The asterisk indicates a significant difference between groups by Welch’s t-test, where *P<0.05. **(C)** Dose response of complement C3 deposition of LA-2 and LA-2 LALAPG in the context of OspA.

To assess the capacity of LA-2 and LA-2 LALAPG to promote complement-dependent borreliacidal activity, we employed a recently developed fluorescence-based *B. burgdorferi* reporter strain GGW979 ^31^. GGW979 is a derivative of *B. burgdorferi* B31 that expresses the red fluorescent protein, mScarlet, under control of an IPTG-inducible promoter ^45^. In the assay, neither LA-2 nor LA-2 LALAPG had any measurable demonstrable borreliacidal activity in the absence of 5 nM complement (**Figure 3**). In the presence of exogenous complement, LA-2 IgG elicited dose-dependent borreliacidal activity at concentrations ranging from 20 to <1 μg/ml (**data not shown**). LA-2 LALAPG, on the other hand, had no measurable complement-dependent borreliacidal activity, even at 20 μg/ml. Collectively, these results demonstrate that the LA-2 LALAPG retains OspA binding activity but lacks complement-fixing activity.

**Figure 3.**
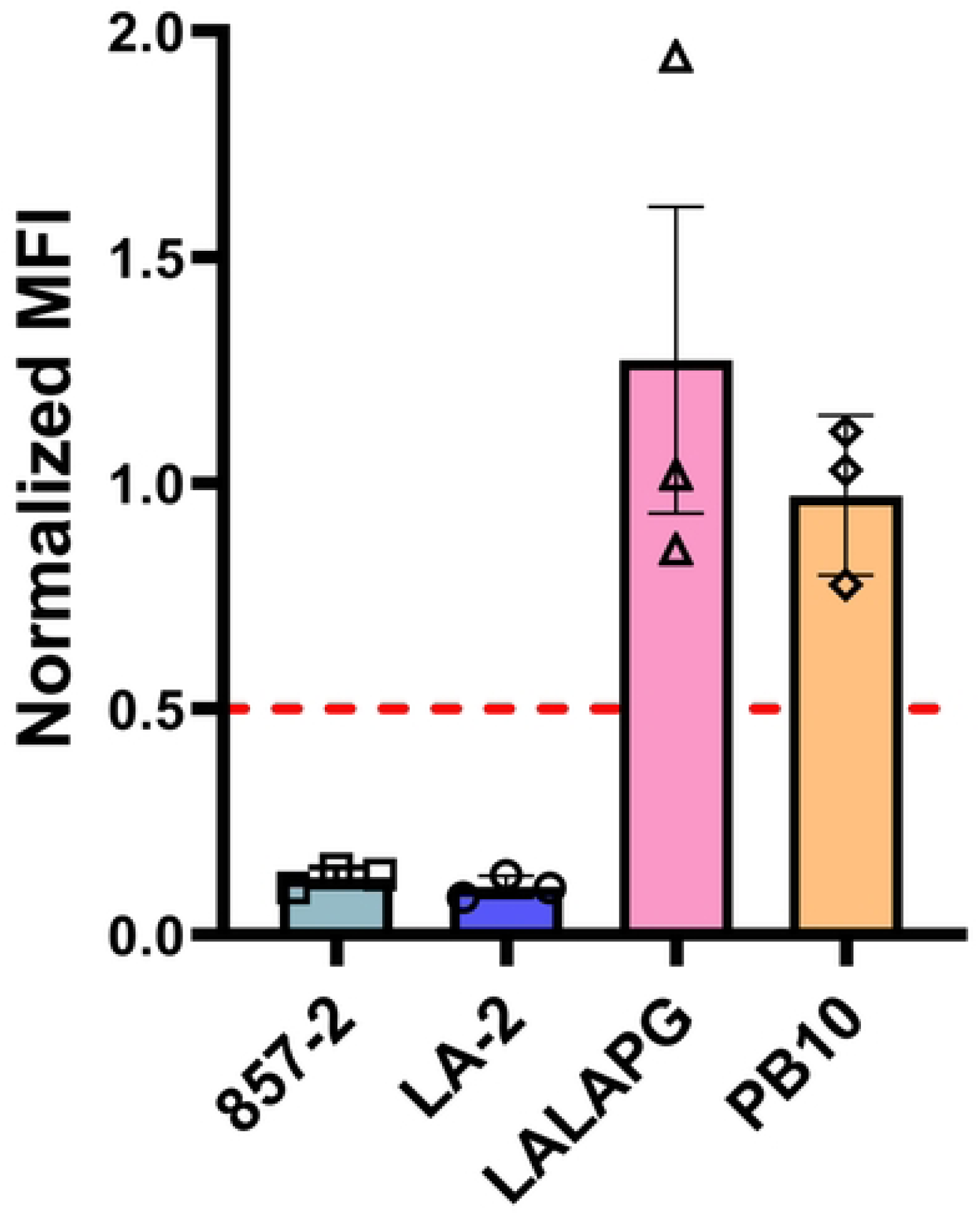
Complement-dependent borreliacidal activity associated with LA-2 and LA-2 LALAPG. Mid-log phase *B. burgdorferi* strain B31-5A4 carrying an IPTG-inducible *mscarlet-I* reporter (GGW979) were suspended (1.5×10⁶ cells per reaction) in BSKII medium supplemented with 20% guinea pig complement and 5 nM of the mAbs indicated on the x-axis (857-2, LA-2, LA-2 LALAPG, PB10), as detailed in the Materials and Methods. Following a 48-h incubation, the median fluorescence intensity (MFI; 569 nm excitation/611 nm emission) was determined. The bars are the mean of three independent experiments with each symbol being an independent experiment and the error bars indicating SD. The dashed red line represented 50% killing. The results demonstrate that 857-2 and LA-2 have potent borreliacidal activity as indicated by low normalized MFI, whereas LA-2 LALAPG and the isotype control were devoid of activity.

### LA-2 LALAPG protects mice from tick-mediated *B. burgdorferi* infection

Having established that LA-2 LALAPG is deficient in complement fixation, we next examined the mAb’s ability to protect mice from infection in a tick-mediated *B. burgdorferi* challenge. Groups of C3H/HeN mice were administered 120 or 30 μg of LA-2 or LA-2 LALAPG by subcutaneous injection and challenged the following day with *B. burgdorferi* B31-infected *Ixodes scapularis* nymphs. An additional group of mice received an IgG1 isotype control (ZAC-3). On day 21, the mice were euthanized and assessed for *B. burgdorferi* infection by serology using a *B.burgdorferi* specific MIA and recovery of viable spirochetes from bladders. For statistical purposes, a mouse was scored as categorically infected if either readout (seroconversion, culture) was positive. By these metrics, LA-2 and LA-2 LALAPG were each protective at the 120 μg dose (p<0.01), but only marginally effective at 30 μg dose, relative to mice that received the isotype control (**Table 2**). Of particular importance, LA-2 and LA-2 LALAPG were statistically indistinguishable in terms of their protective efficacy (p>0.99). These results demonstrate that LA-2 can protect mice from tick-mediated *B. burgdorferi* infection in the absence of complement fixation.

**Table 2.** mAb passive protection in mouse model of tick-mediated *B. burgdorferi* challenge.

| mAb | Dose (µg) | Tick | readout (# pos./# total) |  | significance (p)<sup>b</sup> |  |
| --- | --- | --- | --- | --- | --- | --- |
|  |  |  | serology<sup>a</sup> | culture<sup>a</sup> | vs. IC | vs. LA-2 |
| LA-2 | 120 | + | 0/6 | 0/6 | <0.01 | - |
| LALAPG | 120 | + | 1/6 | 1/6 | 0.03 | >0.99 |
| LA-2 | 30 | + | 2/5 | 1/5 | 0.18 | - |
| LALAPG | 30 | + | 3/6 | 2/6 | 0.18 | >0.99 |
| IC | 30 | + | 5/5 | 5/5 | - | - |
| IC | 30 | - | 0/2 | 0/2 | - | - |

<sup>a</sup>, number of positive mice/total mice per group. <sup>b</sup>, significance (Fisher’s exact test with Benjamani-Hochberg procedure for the FDR) was determined using readout with highest infection status. p-values <0.05 are considered significant. IC = isotype control.

*B. burgdorferi* numbers in the midgut of ticks that feed on OspA immunized mice were reported to decline or be eliminated entirely within days after engorgement suggesting that OspA antibodies exhibit borreliacidal activities within the tick gut ^9,32,46^. On the other hand, the “LA-2-like” mAb, C3.78, passively protected mice from tick-mediated *B. burgdorferi* infection without affecting spirochete numbers in the midgut, consistent with a mechanism of action not dependent on borreliacidal activity ^47^. To address this issue in the case of LA-2, we collected engorged ticks that had fed on ZAC-3-, LA-2- or LA-2 LALAPG-treated mice, dissected the midguts, then quantified spirochete burdens using qPCR. There was no significant reduction in spirochete burdens in the tick midgut in ticks that fed on LA-2 or LA-2 LALAPG-treated mice (**Figure S2**), as compared to the isotype control-treated group. Thus, LA-2 or LA-2 LALAPG do not appear to exhibit borreliacidal activity in the context of the tick midgut environment.

### LA-2 and LA-2 LALAPG limit skin dissemination of *B. burgdorferi*

While it is known that LA-2 and other OspA antibodies protect mice from *B. burgdorferi* dissemination even when spirochetes are delivered by injection, there are no reports examining whether this is dependent on complement ^6,48,49^. Considering the sensitivity of *B. burgdorferi* to the classical complement pathway ^50^, we reasoned that LA-2 would prevent disseminated infection following intradermal challenge, while LA-2 LALAPG would not. To test this, groups of mice were administered LA-2 or LA-2 LALAPG at 120 μg per mouse then challenged the following day with viable *B. burgdorferi* B31 (10^5^ cells) by intradermal injection. Three weeks later, mice were euthanized and assessed for seroconversion using the *B. burgdorferi*-specific MIA, described above. By this measure, all six mice in the LA-2 treated group and five of the six mice in the LA-2 LALAPG group were protected (**Table 3**). This demonstrates that neither Fc effector functions nor complement fixing activity are required for LA-2’s protective activity following intradermal challenge.

**Table 3.**
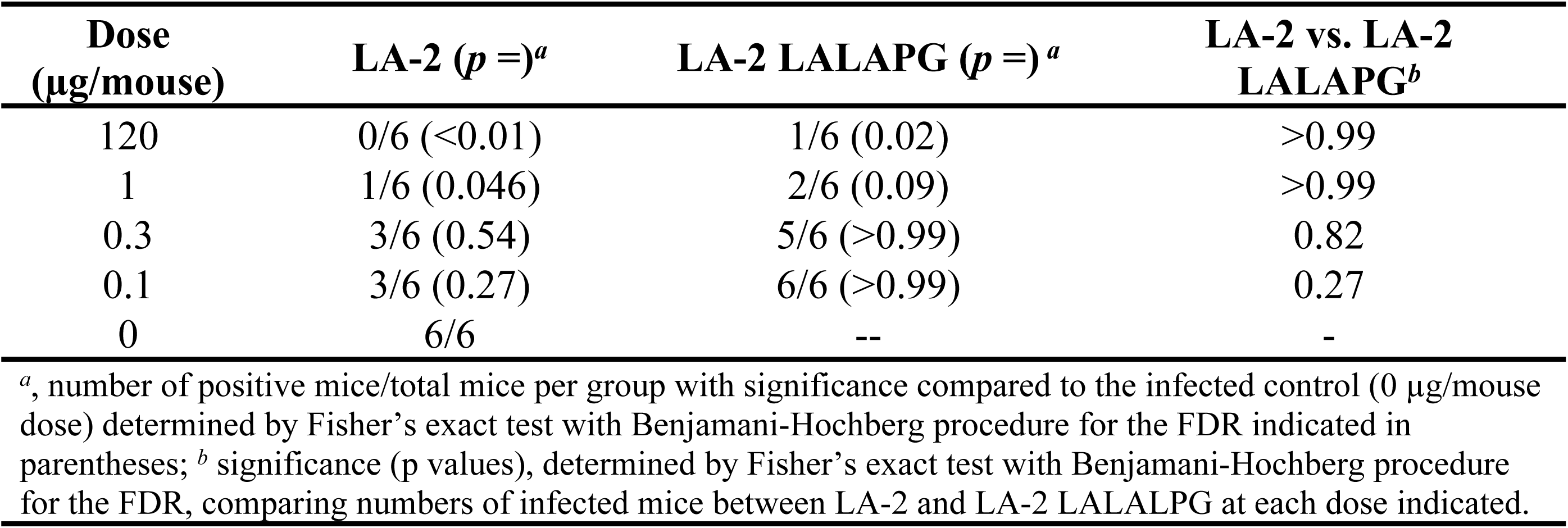
mAb passive protection in mouse model of *B. burgdorferi* ID challenge.

To investigate how LA-2 and LA-2 LALAPG perform at limiting doses, we carried out a pilot study to establish the minimum amount of LA-2 required to protect BALB/c mice against *B. burgdorferi* B31 intradermal challenge. Those studies indicated that as little as 1 μg of LA-2 IgG per mouse was sufficient to render *B. burgdorferi* B31 non-infectious (**Table S1**). We therefore compared LA-2 and LA-2 LALAPG side by side at doses of 1, 0.3 and 0.1 μg mAb per animal in intradermal *B. burgdorferi* B31 challenge. At the 1 μg dose, one of the six mice in the LA-2 treatment group was infected at day 21, while two of the six mice in the LA-2 LALAPG group were infected (**Table 3**). Although neither mAb conferred significant protection at the two lower doses tested (0.3 and 0.1 μg per mouse), LA-2 treated mice fared slightly better than the LA-2 LALAPG treated mice in both cases. We conclude that LA-2 IgG protection in the intradermal challenge model is independent of Fc-mediated activities at high antibody concentrations but possibly important when antibody is limiting.

### LA-2 and LA-2 LALAPG clear viable spirochetes from the *B. burgdorferi* skin

The fact that both LA-2 and LA-2 LALAPG treatments inhibited *B. burgdorferi* dissemination in the mouse model of intradermal challenge prompted us to examine spirochete burden in tissues at earlier time points. To do this, skin biopsies were collected on days 1, 3 and 7 at three locations: the injection site (IS) on the ventral side of the animal, ∼1 cm from the IS, and ∼3 cm from the IS on the dorsal side of the animal (**Figure 4**). Skin biopsies were cultured in BSKII medium for a month and scored regularly for appearance of viable spirochetes. In the control mice, viable spirochetes were recovered on days 1, 3 and 7, which coincides with the reported kinetics of *B. burgdorferi* dissemination ^51,52^. Moreover, three of the four mice also had positive knee and spleen cultures (**data not shown**). In contrast, skin biopsies from LA-2-treated mice were culture negative at all time points examined (**Figure 4**). The results were similar for LA-2 LALAPG treatment, with just one of three animals showing positive cultures on day 7 (**Figure 4**). These results indicate that, in the presence of LA-2 and LA-2 LALAPG, viable *B. burgdorferi* spirochetes are cleared rapidly at or very near the injection site, thereby arresting dissemination before it even gets started.

**Figure 4.**
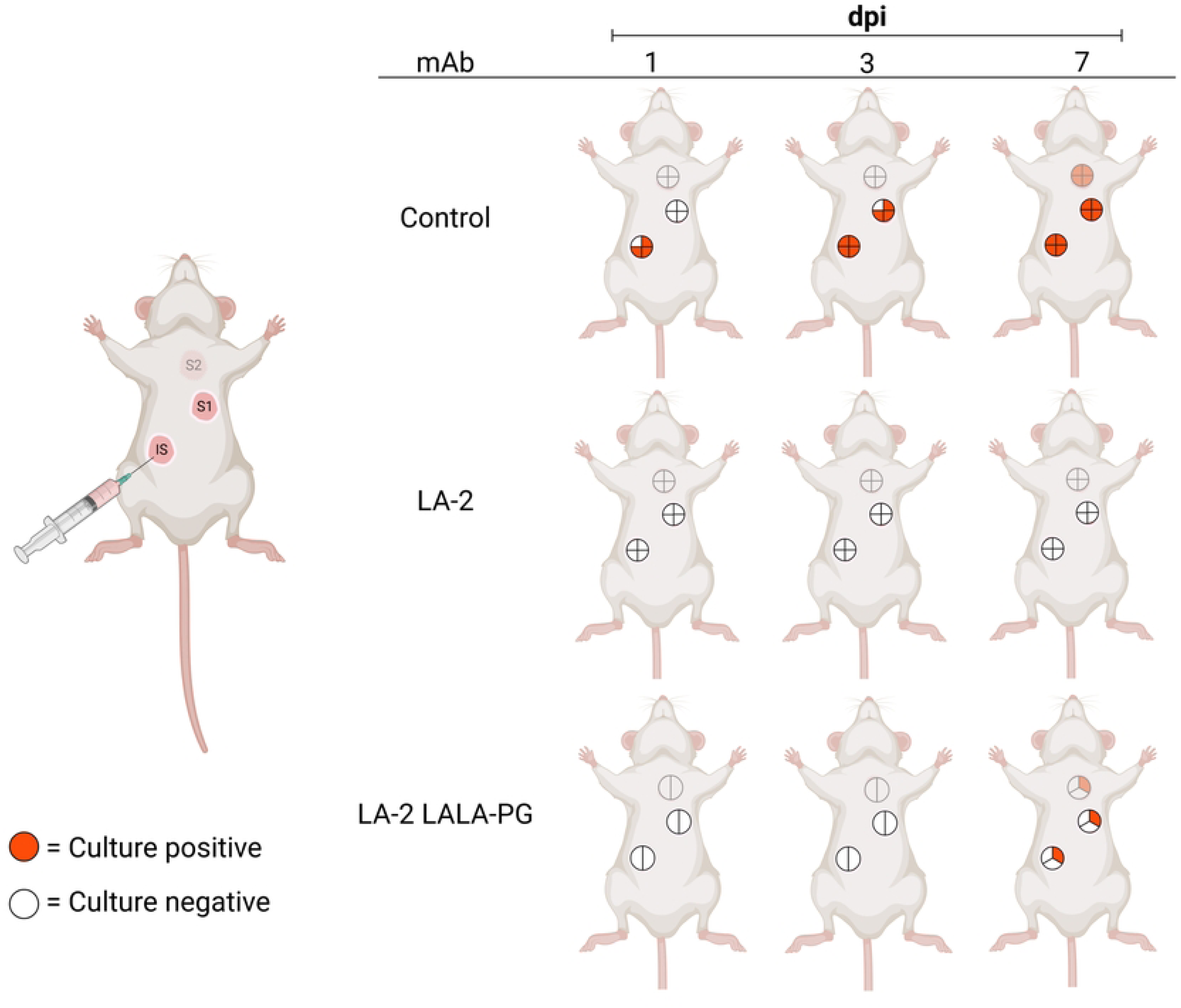
LA-2 and LA-2 LALAPG prevent spirochete dissemination through skin. Groups of mice were administered LA-2, LA-2 LALAPG or an isotype control by subcutaneous injection, then intradermally challenged one day later with *B. burgdorferi.* On days 1, 3, and 7 post challenge, mice were euthanized, and skin biopsies were harvested at the injection site (IS), ∼ 1 cm from the IS (S1), and ∼3 cm from the IS (S2). Biopsies were immersed in BSKII media to recover viable spirochetes. The pie charts indicate the number of skin samples assayed per treatment, with one sample collected from each skin site per mouse. Red subdivisions indicate positivity for motile spirochetes, while white subdivisions indicate no viable spirochetes detected. Shown are the combined results from two independent experiments.

The absence of viable spirochetes in the skin biopsies led us to hypothesize that local inflammation may contribute to LA-2-mediated spirochete clearance. To test this, we examined skin biopsy homogenates for the presence of mouse inflammatory chemokines and chemokines TNF-α, IFN-γ, MCP-1, IL-6, IL-10, and IL-12p70. At five days following injection, we observed elevated levels of TNF-α, IFN-γ, IL-6 and especially MCP-1 in untreated mice when compared to uninfected mice (**Figure 5**). However, in infected mice that were pretreated with LA-2, analytes showed cytokine concentrations similar to levels of uninfected mice (**Figure 5**). Thus, LA-2 treatment is not associated with residual inflammation in the skin and may clear the spirochetes before a cytokine response can be generated. Further analysis of mouse skin cytokines and chemokines, as well as immune cell infiltrates, collected at earlier infection time points is ongoing.

**Figure 5.**
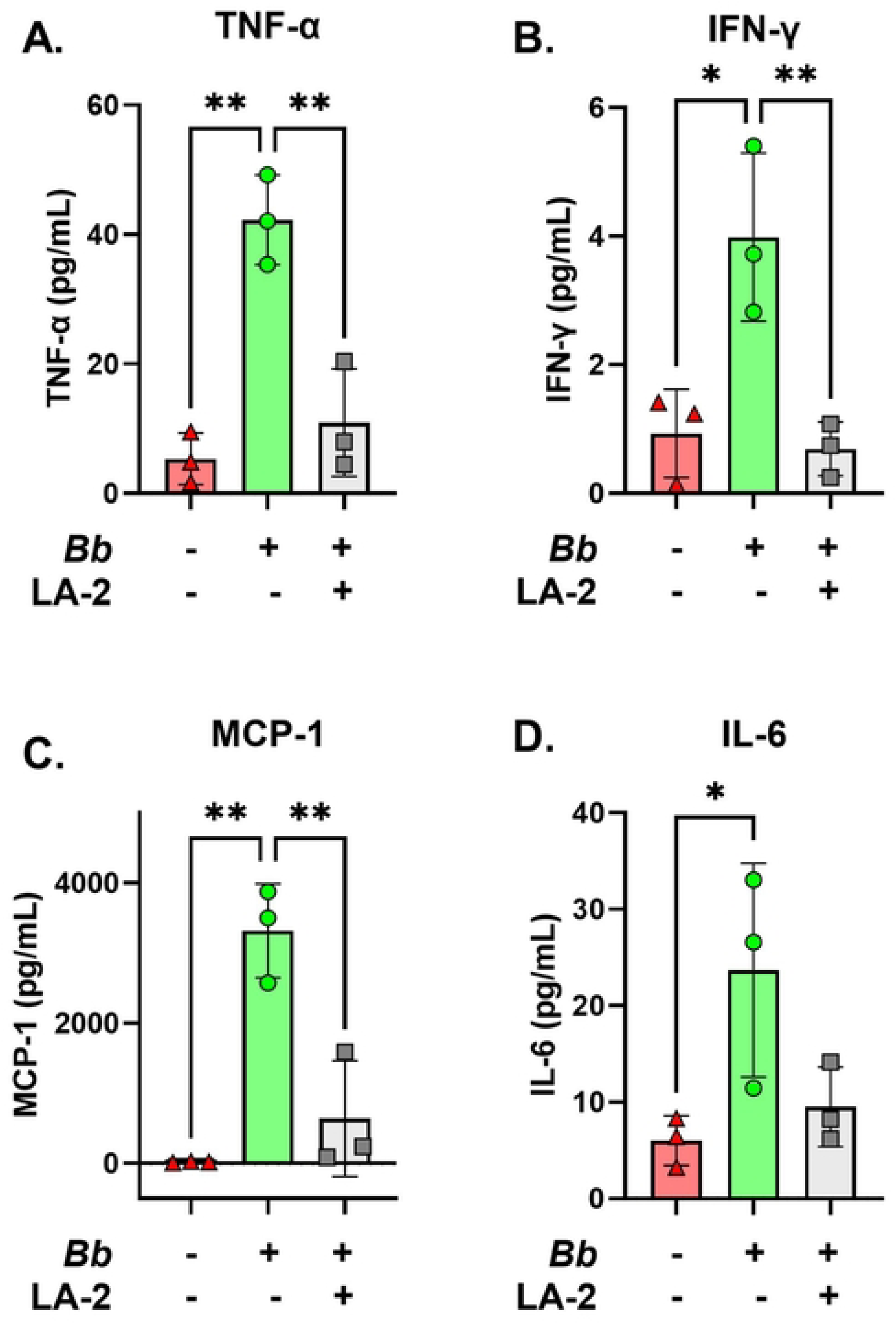
BALB/c mice intradermal infected with *B. burgdorferi* and treated with LA-2, 5 days post-injection. On day –1, mice were given a subcutaneous injection of 30 µg/mL of LA-2 in 200 µL of 1X PBS, 30 µg/mL of ZAC-3 in 200 µL of 1X PBS, or 200 µL of 1X PBS. Mice were shaved on day 0 on their right and left flanks, then given an intradermal injection, on each flank, of 10^5 live *B. burgdorferi* cells in 50 µL of 1X PBS, or 50 µL of 1X PBS. Skin biopsies, approximately 1 cm in diameter, were collected in 1 mL of cytokine extraction buffer for cytometric bead array (CBA) analysis, 5 days post-injection. Skin samples were diluted 1:2 in assay diluent for analysis. Statistics performed by one-way ANOVA, no matching or pairing, corrected for multiple comparisons using Tukey’s test. 95% confidence interval. *p<0.05, **p<0.01.

## Discussion

In this report, we generated and characterized an “Fc-silent” derivative of LA-2 as a tool to investigate the role of complement in passive protection afforded by LA-2 in both tick- and needle-mediated *B. burgdorferi* challenge models. The Fc element of LA-2 was rendered silent by the addition of the so-called LALAPG substitutions (L234A, L235A, P329G), a modification that is gaining wide recognition for its research and clinical applications ^44,53,54^. We confirmed that LA-2 LALAPG retained OspA binding activity comparable to the parenteral LA-2 IgG1 but was markedly attenuated for *in vitro* complement fixation and complement-dependent borreliacidal activity.

When tested *in vivo*, we found that LA-2 LALAPG was as effective as LA-2 IgG in passively protecting mice from tick-mediated *B. burgdorferi* challenge, indicating that neither antibody-mediated complement fixation nor complement-dependent borreliacidal activity were necessary to inhibit spirochete infectivity. In this respect, our results agree with Gipson and de Silva who reported that the “LA-2-like” monoclonal antibody C3.78 blocks tick transmission of *B. burgdorferi* in the absence of host complement ^8,47^. Those studies were conducted using complement-deficient (C3) mice and C3.78 Fab fragments. de Silva and colleagues also demonstrated that C3-deficient mice actively immunized with OspA were also protected against *B. burgdorferi* infection, further reinforcing the notion that host complement is not required for transmission blocking activity of OspA vaccines ^32^.

Furthermore, our results support a model in which LA-2 and LA-2 LALAPG inhibit *B. burgdorferi* transmission without affecting the number of spirochetes within the tick midgut. This observation is consistent with C3.78’s mode of action, in which low dose (∼60 μg/mouse) antibody protected mice from tick-mediated *B. burgdorferi* challenge without a concomitant reduction in spirochete numbers in tick tissues ^47^. In other words, antibody is proposed to block spirochete egress from the midgut by a non-borreliacidal mechanism. Gipson and de Silva and others have speculated that OspA antibodies like C3.78 influence the expression of spirochete genes and gene products required for transmission, including the requisite OspA to OspC transition ^23,47^. We favor a model in which OspA antibodies like LA-2 entrap spirochetes within the midgut by physically altering their transmigratory activity ^34^. Using a two compartment Transwell system, we reported recently that spirochete movement from the lower to upper chambers is reduced by >99% in the presence LA-2 or LA-2 LALAPG. Inhibition of transmigration coincided with LA-2’s ability to promote spirochete agglutination, alternations in membrane permeability, and even bleb formation ^33,55^. Exactly how LA-2 engagement with OspA results in changes in migratory activity remains obscure.

While LA-2 has the capacity to interfere with *B. burgdorferi* transmission within the context of the tick, it can also reduce infectivity of *B. burgdorferi* within the mouse. Indeed, LA-2 was originally identified as being capable of passively protecting *scid* mice from subcutaneous *B. burgdorferi* challenge ^6^. We confirmed and extended that original observation by demonstrating in both BALB/c and C3H mice that remarkably low doses of passively administered LA-2 were sufficient to not only confine but seemingly eliminate *B. burgdorferi* from the site of intradermal inoculation within hours. LA-2 LALAPG had similar properties, indicating that clearance of *B. burgdorferi* from the skin environment occurs without complement or Fc effector functions. These observations may be of clinical importance, as they suggest that if *B. burgdorferi* evades immunity within the context of the tick body, any spirochetes that still express OspA upon entry into the skin will encounter a second line of defense ^21,56^. While the underlying mechanism by which LA-2 promotes clearance of spirochetes from the skin environment without Fc effector functions is unknown, there are interesting parallels with antibody-mediated clearance of malaria parasites in this same environment that involve motility arrest and membrane shedding ^57,58^.

In summary, we have demonstrated that LA-2, the well-characterized monoclonal antibody directed against the C-terminus of OspA, functions in both the tick and mammalian environments to limit *B. burgdorferi* infection without the need for Fc effector functions, such as complement fixation and FcγR interactions. It is unclear whether LA-2 equivalence, as defined by a competitive ELISA, which correlates with immunity to Lyme disease in animal models and humans reflects functional activities in vivo or simply a proxy for other activities. Nonetheless, our study makes a case for LA-2’s primary mode of action involving direct physical interactions with the spirochete rather than complement-dependent killing. Elucidating these mechanisms may have implications for our understanding of the mechanistic correlates of OspA-based vaccine-induced immunity in humans.

## Acknowledgements

We are grateful to Dr. Michael Pauly and colleagues ZabBio for generating LA-2 LALAPG. We thank Drs. Renji Song and Jennifer Yates of the Wadsworth Center’s Immunology Core for assistance with flow cytometry and the Media and Cell Culture core for BSK II medium. We thank Ms. Elizabeth Cavosie (Wadsworth Center) for administrative assistance. BioRender was used for some figure generation. This work was supported by the National Institute of Allergy and Infectious Diseases (NIAID), National Institutes of Health, Department of Health and Human Services, Contract No. 75N93019C00040 (PI/PD Mantis). This content is solely the responsibility of the authors and does not necessarily represent the official views of the NIH.

## References

1. Lanzavecchia, A., Fruhwirth, A., Perez, L., and Corti, D. (2016). Antibody-guided vaccine design: identification of protective epitopes. Curr Opin Immunol 41, 62–67. 10.1016/j.coi.2016.06.001.

2. Principato, S., Pizza, M., and Rappuoli, R. (2020). Meningococcal factor H binding protein as immune evasion factor and vaccine antigen. FEBS Lett 594, 2657–2669. 10.1002/1873-3468.13793.

3. Yoo, R., Jore, M.M., and Julien, J.P. (2025). Targeting Bottlenecks in Malaria Transmission: Antibody-Epitope Descriptions Guide the Design of Next-Generation Biomedical Interventions. Immunol Rev 330, e70001. 10.1111/imr.70001.

4. Fikrig, E., Barthold, S.W., Kantor, F.S., and Flavell, R.A. (1990). Protection of mice against the Lyme disease agent by immunizing with recombinant OspA. Science 250, 553–556. 10.1126/science.2237407.

5. Kramer, M.D., Schaible, U.E., Wallich, R., Moter, S.E., Petzoldt, D., and Simon, M.M. (1990). Characterization of Borrelia burgdorferi associated antigens by monoclonal antibodies. Immunobiology 181, 357–366. 10.1016/s0171-2985(11)80504-8.

6. Schaible, U.E., Kramer, M.D., Eichmann, K., Modolell, M., Museteanu, C., and Simon, M.M. (1990). Monoclonal antibodies specific for the outer surface protein A (OspA) of Borrelia burgdorferi prevent Lyme borreliosis in severe combined immunodeficiency (scid) mice. Proc Natl Acad Sci U S A 87, 3768–3772. 10.1073/pnas.87.10.3768.

7. Fikrig, E., Barthold, S.W., Marcantonio, N., Deponte, K., Kantor, F.S., and Flavell, R.A. (1992). Roles of OspA, OspB, and flagellin in protective immunity to Lyme borreliosis in laboratory mice. Infect Immun 60, 657–661. 10.1128/iai.60.2.657-661.1992.

8. Sears, J.E., Fikrig, E., Nakagawa, T.Y., Deponte, K., Marcantonio, N., Kantor, F.S., and Flavell, R.A. (1991). Molecular mapping of Osp-A mediated immunity against Borrelia burgdorferi, the agent of Lyme disease. J Immunol 147, 1995–2000.

9. de Silva, A.M., Telford, S.R., 3rd, Brunet, L.R., Barthold, S.W., and Fikrig, E. (1996). Borrelia burgdorferi OspA is an arthropod-specific transmission-blocking Lyme disease vaccine. J Exp Med 183, 271–275.

10. Sădziene, A., Rosa, P.A., Thompson, P.A., Hogan, D.M., and Barbour, A.G. (1992). Antibody-resistant mutants of Borrelia burgdorferi: in vitro selection and characterization. J Exp Med 176, 799–809. 10.1084/jem.176.3.799.

11. Sadziene, A., Thompson, P.A., and Barbour, A.G. (1993). In vitro inhibition of Borrelia burgdorferi growth by antibodies. J Infect Dis 167, 165–172. 10.1093/infdis/167.1.165.

12. Sadziene, A., Jonsson, M., Bergström, S., Bright, R.K., Kennedy, R.C., and Barbour, A.G. (1994). A bactericidal antibody to Borrelia burgdorferi is directed against a variable region of the OspB protein. Infect Immun 62, 2037–2045. 10.1128/iai.62.5.2037-2045.1994.

13. Wilske, B., Luft, B., Schubach, W.H., Zumstein, G., Jauris, S., Preac-Mursic, V., and Kramer, M.D. (1992). Molecular analysis of the outer surface protein A (OspA) of Borrelia burgdorferi for conserved and variable antibody binding domains. Med Microbiol Immunol 181, 191–207. 10.1007/bf00215765.

14. Wilske, B., Preac-Mursic, V., Gobel, U.B., Graf, B., Jauris, S., Soutschek, E., Schwab, E., and Zumstein, G. (1993). An OspA serotyping system for Borrelia burgdorferi based on reactivity with monoclonal antibodies and OspA sequence analysis. J Clin Microbiol 31, 340–350. 10.1128/jcm.31.2.340-350.1993.

15. LaRocca, T.J., and Benach, J.L. (2008). The important and diverse roles of antibodies in the host response to Borrelia infections. Curr Top Microbiol Immunol 319, 63–103.

16. Pal, U., de Silva, A.M., Montgomery, R.R., Fish, D., Anguita, J., Anderson, J.F., Lobet, Y., and Fikrig, E. (2000). Attachment of Borrelia burgdorferi within Ixodes scapularis mediated by outer surface protein A. J Clin Invest 106, 561–569. 10.1172/JCI9427.

17. Li, H., and Lawson, C.L. (1995). Crystallization and preliminary X-ray analysis of Borrelia burgdorferi outer surface protein A (OspA) complexed with a murine monoclonal antibody Fab fragment. J Struct Biol 115, 335–337. 10.1006/jsbi.1995.1058.

18. Ding, W., Huang, X., Yang, X., Dunn, J.J., Luft, B.J., Koide, S., and Lawson, C.L. (2000). Structural identification of a key protective B-cell epitope in Lyme disease antigen OspA. J Mol Biol 302, 1153–1164. 10.1006/jmbi.2000.4119.

19. Li, H., Dunn, J.J., Luft, B.J., and Lawson, C.L. (1997). Crystal structure of Lyme disease antigen outer surface protein A complexed with an Fab. Proc Natl Acad Sci U S A 94, 3584–3589.

20. Schubach, W.H., Mudri, S., Dattwyler, R.J., and Luft, B.J. (1991). Mapping antibody-binding domains of the major outer surface membrane protein (OspA) of Borrelia burgdorferi. Infect Immun 59, 1911–1915. 10.1128/iai.59.6.1911-1915.1991.

21. Caimano, M.J., Eggers, C.H., Gonzalez, C.A., and Radolf, J.D. (2005). Alternate sigma factor RpoS is required for the in vivo-specific repression of Borrelia burgdorferi plasmid lp54-borne ospA and lp6.6 genes. J Bacteriol 187, 7845–7852. 10.1128/JB.187.22.7845-7852.2005.

22. Schwan, T.G., Piesman, J., Golde, W.T., Dolan, M.C., and Rosa, P.A. (1995). Induction of an outer surface protein on Borrelia burgdorferi during tick feeding. Proc Natl Acad Sci U S A 92, 2909–2913. 10.1073/pnas.92.7.2909.

23. Wang, Y., Kern, A., Boatright, N.K., Schiller, Z.A., Sadowski, A., Ejemel, M., Souders, C.A., Reimann, K.A., Hu, L., Thomas, W.D., Jr., and Klempner, M.S. (2016). Pre-exposure Prophylaxis With OspA-Specific Human Monoclonal Antibodies Protects Mice Against Tick Transmission of Lyme Disease Spirochetes. J Infect Dis 214, 205–211. 10.1093/infdis/jiw151.

24. Schiller, Z.A., Rudolph, M.J., Toomey, J.R., Ejemel, M., LaRochelle, A., Davis, S.A., Lambert, H.S., Kern, A., Tardo, A.C., Souders, C.A., et al. (2021). Blocking Borrelia burgdorferi transmission from infected ticks to nonhuman primates with a human monoclonal antibody. J Clin Invest 131. 10.1172/JCI144843.

25. Rudolph, M.J., Davis, S.A., Haque, H.M.E., Ejemel, M., Cavacini, L.A., Vance, D.J., Willsey, G.G., Piazza, C.L., Weis, D.D., Wang, Y., and Mantis, N.J. (2023). Structure of a transmission blocking antibody in complex with Outer surface protein A from the Lyme disease spirochete, Borreliella burgdorferi. Proteins. 10.1002/prot.26549.

26. Golde, W.T., Piesman, J., Dolan, M.C., Kramer, M., Hauser, P., Lobet, Y., Capiau, C., Desmons, P., Voet, P., Dearwester, D., and Frantz, J.C. (1997). Reactivity with a specific epitope of outer surface protein A predicts protection from infection with the Lyme disease spirochete, Borrelia burgdorferi. Infect Immun 65, 882–889.

27. Van Hoecke, C., Comberbach, M., De Grave, D., Desmons, P., Fu, D., Hauser, P., Lebacq, E., Lobet, Y., and Voet, P. (1996). Evaluation of the safety, reactogenicity and immunogenicity of three recombinant outer surface protein (OspA) lyme vaccines in healthy adults. Vaccine 14, 1620–1626. 10.1016/s0264-410x(96)00146-6.

28. Steere, A.C., Sikand, V.K., Meurice, F., Parenti, D.L., Fikrig, E., Schoen, R.T., Nowakowski, J., Schmid, C.H., Laukamp, S., Buscarino, C., and Krause, D.S. (1998). Vaccination against Lyme disease with recombinant Borrelia burgdorferi outer-surface lipoprotein A with adjuvant. Lyme Disease Vaccine Study Group. N Engl J Med 339, 209–215. 10.1056/NEJM199807233390401.

29. Tahir, D., Geolier, V., Bruant, H., Le Flèche-Matéos, A., Mallet, A., Varloud, M., Civat, C., Girerd-Chambaz, Y., Montano, S., Pion, C., et al. (2025). A Lyme disease mRNA vaccine targeting Borrelia burgdorferi OspA induces strong immune responses and prevents transmission in mice. Mol Ther Nucleic Acids 36, 102514. 10.1016/j.omtn.2025.102514.

30. Haque, H.M.E., Ejemel, M., Vance, D.J., Willsey, G., Rudolph, M.J., Cavacini, L.A., Wang, Y., Mantis, N.J., and Weis, D.D. (2022). Human B Cell Epitope Map of the Lyme Disease Vaccine Antigen, OspA. ACS Infect Dis. 10.1021/acsinfecdis.2c00346.

31. Vance, D.J., Basir, S., Piazza, C.L., Willsey, G.G., Haque, H.M.E., Tremblay, J.M., Rudolph, M.J., Muriuki, B., Cavacini, L., Weis, D.D., et al. (2024). Single-domain antibodies reveal unique borrelicidal epitopes on the Lyme disease vaccine antigen, outer surface protein A (OspA). Infect Immun 92, e0008424. 10.1128/iai.00084-24.

32. Rathinavelu, S., Broadwater, A., and de Silva, A.M. (2003). Does host complement kill Borrelia burgdorferi within ticks? Infect Immun 71, 822–829. 10.1128/iai.71.2.822-829.2003.

33. Frye, A.M., Ejemel, M., Cavacini, L., Wang, Y., Rudolph, M.J., Song, R., and Mantis, N.J. (2022). Agglutination of Borreliella burgdorferi by Transmission-Blocking OspA Monoclonal Antibodies and Monovalent Fab Fragments. Infect Immun, e0030622. 10.1128/iai.00306-22.

34. Bhattacharyya, A., and Mantis, N.J. (2025). OspA Antibodies Inhibit the In vitro Transmigration of *Borreliellia burgdorferi*. bioRxiv, 2025.2011.2020.689610. 10.1101/2025.11.20.689610.

35. Bézay, N., Wagner, L., Kadlecek, V., Obersriebnig, M., Wressnigg, N., Hochreiter, R., Schneider, M., Dubischar, K., Derhaschnig, U., Klingler, A., et al. (2024). Optimisation of dose level and vaccination schedule for the VLA15 Lyme borreliosis vaccine candidate among healthy adults: two randomised, observer-blind, placebo-controlled, multicentre, phase 2 studies. Lancet Infect Dis 24, 1045–1058. 10.1016/s1473-3099(24)00175-0.

36. Lundberg, U., Hochreiter, R., Timofoyeva, Y., Kanevsky, I., Meinke, A., Anderson, A.S., and Simon, R. (2024). Preclinical Evidence for the Protective Capacity of Antibodies Induced by Lyme Vaccine Candidate VLA15 in People. Open Forum Infect Dis 11, ofae467. 10.1093/ofid/ofae467.

37. Rudolph, M.J., Davis, S.A., Haque, H.M.E., Weis, D.D., Vance, D.J., Piazza, C.L., Ejemel, M., Cavacini, L., Wang, Y., Mbow, M.L., et al. (2023). Structural Elucidation of a Protective B Cell Epitope on Outer Surface Protein C (OspC) of the Lyme Disease Spirochete, Borreliella burgdorferi. mBio 14, e0298122. 10.1128/mbio.02981-22.

38. Rudolph, M.J., Muriuki, B.M., Chen, Y., Vance, D.J., Vorauer, C., Piazza, C.L., Freeman-Gallant, G., Golonka, R.M., Mirabile, G., Guttman, M., et al. (2025). Germline encoded residues dominate the interaction of a human monoclonal antibody with decorin binding protein A of Borrelia burgdorferi. Front Immunol 16, 1611828. 10.3389/fimmu.2025.1611828.

39. Giritch, A., Marillonnet, S., Engler, C., van Eldik, G., Botterman, J., Klimyuk, V., and Gleba, Y. (2006). Rapid high-yield expression of full-size IgG antibodies in plants coinfected with noncompeting viral vectors. Proc Natl Acad Sci U S A 103, 14701–14706. 10.1073/pnas.0606631103.

40. Swope, K., Morton, J., Pogue, G.P., Hume, S., Pauly, M.H., Shepherd, J., Simpson, C.A., Bratcher, B., Whaley, K.J., Zeitlin, L., et al. (2021). Manufacturing plant-made monoclonal antibodies for research or therapeutic applications. Methods Enzymol 660, 239–263. 10.1016/bs.mie.2021.05.011.

41. Fischinger, S., Fallon, J.K., Michell, A.R., Broge, T., Suscovich, T.J., Streeck, H., and Alter, G. (2019). A high-throughput, bead-based, antigen-specific assay to assess the ability of antibodies to induce complement activation. J Immunol Methods 473, 112630. 10.1016/j.jim.2019.07.002.

42. Kocan, K.M., de la Fuente, J., and Coburn, L.A. (2015). Insights into the development of Ixodes scapularis: a resource for research on a medically important tick species. Parasit Vectors 8, 592. 10.1186/s13071-015-1185-7.

43. Schlothauer, T., Herter, S., Koller, C.F., Grau-Richards, S., Steinhart, V., Spick, C., Kubbies, M., Klein, C., Umaña, P., and Mössner, E. (2016). Novel human IgG1 and IgG4 Fc-engineered antibodies with completely abolished immune effector functions. Protein Eng Des Sel 29, 457–466. 10.1093/protein/gzw040.

44. Wilkinson, I., Anderson, S., Fry, J., Julien, L.A., Neville, D., Qureshi, O., Watts, G., and Hale, G. (2021). Fc-engineered antibodies with immune effector functions completely abolished. PLoS One 16, e0260954. 10.1371/journal.pone.0260954.

45. Bindels, D.S., Haarbosch, L., van Weeren, L., Postma, M., Wiese, K.E., Mastop, M., Aumonier, S., Gotthard, G., Royant, A., Hink, M.A., and Gadella, T.W., Jr. (2017). mScarlet: a bright monomeric red fluorescent protein for cellular imaging. Nat Methods 14, 53–56. 10.1038/nmeth.4074.

46. Fikrig, E., Telford, S.R., 3rd, Barthold, S.W., Kantor, F.S., Spielman, A., and Flavell, R.A. (1992). Elimination of Borrelia burgdorferi from vector ticks feeding on OspA-immunized mice. Proc Natl Acad Sci U S A 89, 5418–5421.

47. Gipson, C.L., and de Silva, A.M. (2005). Interactions of OspA monoclonal antibody C3.78 with Borrelia burgdorferi within ticks. Infect Immun 73, 1644–1647. 10.1128/IAI.73.3.1644-1647.2005.

48. Simon, M.M., Schaible, U.E., Kramer, M.D., Eckerskorn, C., Museteanu, C., Müller-Hermelink, H.K., and Wallich, R. (1991). Recombinant outer surface protein a from Borrelia burgdorferi induces antibodies protective against spirochetal infection in mice. J Infect Dis 164, 123–132. 10.1093/infdis/164.1.123.

49. Pine, M., Arora, G., Hart, T.M., Bettini, E., Gaudette, B.T., Muramatsu, H., Tombacz, I., Kambayashi, T., Tam, Y.K., Brisson, D., et al. (2023). Development of an mRNA-lipid nanoparticle vaccine against Lyme disease. Mol Ther 31, 2702–2714. 10.1016/j.ymthe.2023.07.022.

50. Kochi, S.K., and Johnson, R.C. (1988). Role of immunoglobulin G in killing of Borrelia burgdorferi by the classical complement pathway. Infect Immun 56, 314–321.

51. Barthold, S.W., Persing, D.H., Armstrong, A.L., and Peeples, R.A. (1991). Kinetics of Borrelia burgdorferi dissemination and evolution of disease after intradermal inoculation of mice. Am J Pathol 139, 263–273.

52. Shih, C.M., Pollack, R.J., Telford, S.R., 3rd, and Spielman, A. (1992). Delayed dissemination of Lyme disease spirochetes from the site of deposition in the skin of mice. J Infect Dis 166, 827–831. 10.1093/infdis/166.4.827.

53. Damelang, T., Brinkhaus, M., van Osch, T.L.J., Schuurman, J., Labrijn, A.F., Rispens, T., and Vidarsson, G. (2023). Impact of structural modifications of IgG antibodies on effector functions. Front Immunol 14, 1304365. 10.3389/fimmu.2023.1304365.

54. Johnson, N.V., Wall, S.C., Kramer, K.J., Holt, C.M., Periasamy, S., Richardson, S.I., Manamela, N.P., Suryadevara, N., Andreano, E., Paciello, I., et al. (2024). Discovery and characterization of a pan-betacoronavirus S2-binding antibody. Structure 32, 1893–1909.e1811. 10.1016/j.str.2024.08.022.

55. Kudryashev, M., Cyrklaff, M., Baumeister, W., Simon, M.M., Wallich, R., and Frischknecht, F. (2009). Comparative cryo-electron tomography of pathogenic Lyme disease spirochetes. Mol Microbiol 71, 1415–1434. 10.1111/j.1365-2958.2009.06613.x.

56. Caimano, M.J., Groshong, A.M., Belperron, A., Mao, J., Hawley, K.L., Luthra, A., Graham, D.E., Earnhart, C.G., Marconi, R.T., Bockenstedt, L.K., et al. (2019). The RpoS Gatekeeper in Borrelia burgdorferi: An Invariant Regulatory Scheme That Promotes Spirochete Persistence in Reservoir Hosts and Niche Diversity. Front Microbiol 10, 1923. 10.3389/fmicb.2019.01923.

57. Vanderberg, J.P., and Frevert, U. (2004). Intravital microscopy demonstrating antibody-mediated immobilisation of Plasmodium berghei sporozoites injected into skin by mosquitoes. Int J Parasitol 34, 991–996. 10.1016/j.ijpara.2004.05.005.

58. Flores-Garcia, Y., Nasir, G., Hopp, C.S., Munoz, C., Balaban, A.E., Zavala, F., and Sinnis, P. (2018). Antibody-Mediated Protection against Plasmodium Sporozoites Begins at the Dermal Inoculation Site. mBio 9. 10.1128/mBio.02194-18.

